# Pencil Beam Based X-ray Luminescence Computed Tomography Imaging of Deep Targets Three-dimensionally and Quantitatively

**DOI:** 10.1101/2024.05.28.596355

**Authors:** Yile Fang, Yibing Zhang, Changqing Li

**Affiliations:** Department of Bioengineering, University of California, Merced, Merced, CA 95343; Department of Electrical Engineering, University of California, Merced, Merced, CA 95343

## Abstract

X-ray luminescence computed tomography (XLCT), especially pencil beam based XLCT, has been emerged for more than 15 years because it can image deep targets in tissue with high spatial resolution and high measurement sensitivity. However, its application is limited by long scanning time and the tedious efforts of finite element mesh generation for model based reconstruction. Our previous study has addressed the imaging speed issue. In this study, we propose to reconstruct XLCT with weighted filtered back projection (wFBP) which makes it possible to perform XLCT imaging three dimensionally for complex objects at a high spatial resolution. We have also proposed measurements with multiple detectors to obtain quantitative XLCT images. To evaluate the proposed methods, we have performed numerical simulations and different sets of phantom experiments including phantom experiments with a control contrast target, different concentrations, and different target sizes. We have also successfully reconstructed deep targets with a diameter of approximately 0.15 mm. For the phantom experiments with targets of different concentrations, using a 4-channel XLCT system and the wFBP, we achieved the ratio of 3.8:1.79:1, closely resembling the ground truth ratio of 4:2:1. Lastly, we have, for the first time, reconstructed different 3D targets at high spatial resolution, successfully, including a printed M target, a triangular hollow bar, and a hot background cuboid with “UCM Li Lab” slots. Our results indicate that the proposed wFBP advances the promising XLCT technology and makes it ready for in vivo studies in the future.

## 1. Introduction

X-ray luminescence computed tomography, XLCT, has been emerged for more than 15 years as a novel hybrid imaging modality with the promising to combine the high spatial resolution of x-ray imaging and high sensitivity of optical imaging [1][2][3][4][5]. In XLCT imaging, the high energy x-ray photons excite x-ray excitable particles such as Gadolinium oxysulfide doped with europium (Gd_2_O_2_S: Eu^3+^) which emit optical photons to be measured for reconstruction [6][7]. There are two major advantages of x-ray excitation. The first one is the removal of autofluorescence in the laser excited fluorescence imaging. The last one is that the high spatial resolution of x-ray imaging can be used as anatomical guidance in XLCT imaging so that XLCT could overcome the optical scattering effects when imaging deep targets in biological tissues. Furthermore, the emission wavelength of the particle is in the near infrared (NIR) range. Thus, the emission optical photons have high penetration power in tissue so that the optical measurement sensitivity is supposed to be high.

Several types of X-ray Luminescence Computed Tomography (XLCT) systems have been designed and investigated so far. Pratx et al. [3] were pioneers in introducing the initial prototype system, laying the foundation for the concept of hybrid X-ray luminescence optical tomography. Li et al.[8] conducted experimental demonstrations showcasing XLCT’s ability to achieve high spatial resolution through collimated X-ray beams for deep-seated targets. In a different approach, Zhang et al.[9] proposed a multiple pinhole-based XLCT design, employing multiple X-ray beams to simultaneously scan objects and thereby reducing data acquisition time, achieving an impressive edge-to-edge distance of 0.6 mm. Chen et al. [4] put forth a cone-beam XLCT design, aiming to improve data acquisition time, albeit at the expense of spatial resolution. Furthermore, Liu et al. [10] applied a cone-beam-based XLCT approach for small animal imaging. Lun et al. [11] reported the successful reconstruction of a phosphor target with a concentration as low as 0.01 mg/mL at a scanning depth of 21 mm. Fang et al. [12] recently reported a superfast X-ray luminescence computed tomography (XLCT) scan scheme utilizing a photon counter detector and fly-scanning method, achieving a remarkable 28.6-fold increase in speed (43 seconds per transverse scan) compared to previous methods, while maintaining improved XLCT image quality. This rapid scanning approach offers a viable method for conducting 3D X-ray Luminescence Computed Tomography (XLCT) imaging. All these findings have illustrated the potential of XLCT as a valuable tool for in vivo small animal imaging.

However, the applications of XLCT are limited by several factors. The first factor is the scanning speed. There are two types of XLCT imaging: cone beam based XLCT [4][13] and pencil beam based XLCT [14][15][16]. The pencil beam XLCT has the ability to utilize the anatomical guidance from the pencil beam of x-ray. However, it takes a very long time to scan objects with the pencil beam based XLCT. We have addressed this problem in our recent paper by introducing a superfast scanning scheme [12]. Another limitation factor is the tedious work for the finite element mesh generation for the model based XLCT reconstruction [17]. It is very hard to generate good finite element mesh for mice with irregular shapes. The last limitation factor is the limited density of the finite element node which makes it hard to obtain super high spatial resolution of the reconstructed XLCT images. In this study, we propose a weighted filtered back projection (wFBP) to address the last two limitation factors.

In the past, most of our studies utilized a single sensitivity optical photon detector [12][14][17][18]. However, the collection efficiency of excited optical photons by a single detector channel is low, consequently limiting the sensitivity and accuracy of the reconstructed XLCT images, while we have demonstrated that it is sufficient to obtain acceptable XLCT images with a single detector. To enhance XLCT imaging, increasing the number of sensitivity optical photon detectors represents a feasible and practical approach. Zhang et al. [16] have recently introduced a multichannel XLCT system with two detectors, achieving successful reconstruction of three targets measuring 0.58 mm each, employing a 150 μm x-ray beam. In this study, we investigate the effects of detector numbers in XLCT imaging numerically and then utilize 4 detectors in the experimental system.

In this paper, we present the benchtop XLCT imaging system with 4 sensitivity optical detectors, the wFBP reconstruction algorithm, the numerical simulation setup, and the phantom geometry in the method section. Then, we describe the results. Finally, we conclude the paper with discussions.

## 2. Methods

### 2.1 Pencil beam based x-ray luminescence computed tomography system with multiple detectors

Based on the previous studies [15][19], we built a pencil beam based XLCT imaging system with four detectors. In the following numerical simulations, we investigated the effect of detector number up to 16 detectors, while in the physical experimental setup, we only utilized 4 detectors due to the limited resource. Fig. 1 shows the schematics of the four-channel XLCT system which was developed from the single-channel XLCT system in our lab [16][15][13]. Fig. 2 displays a photograph of the XLCT imaging system. The x-ray tube, X-Beam Powerflux [Mo anode, XOS] produces x-ray photons with a peak voltage of 50 kVp and a current up to 1 mA. There is an x-ray optics lens used to focus the x-ray beam with a focal spot size of the 150 micrometers (µm) at a focal length of 44.5 mm. The imaged objects such as phantoms were placed on a motorized rotary stage (RT-3, Newmark System Inc) which was 45 mm away from the output of the x-ray optics lens. The rotary stage is mounted on a motorized vertical stage (VS-50, Newmark Systems Inc.), which controls the scanning height of the object. The vertical stage is amount on a motorized linear stage (NLE-100, Newmark Systems Inc.) which moves the object linearly for each angular projection. In this system, we use a piece of crystal (2cm by 2cm) with a liquid lightguide (SERIES 380, Lumatec) and a photomultiplier tube (PMT) (H7422-50, Hamamatsu) detector to sense the pencil beam x-ray intensity as a single pixel detector for the pencil x-ray computed tomography imaging of the object. The PMT signal is amplified using broadband amplifier (SR455A, Stanford Research Systems) with a gain of 125. Then, the signals undergo filtration using a low-pass filter (BLP-10.7+, fc = 11 MHz, Mini-Circuits) to minimize noises. Subsequently, the signals from the filter were acquired by a gated photon counter, SR400 by Stanford Research System. In this study, the single pixel detector is also used to detect the object boundaries during scanning.

**Figure 1.**
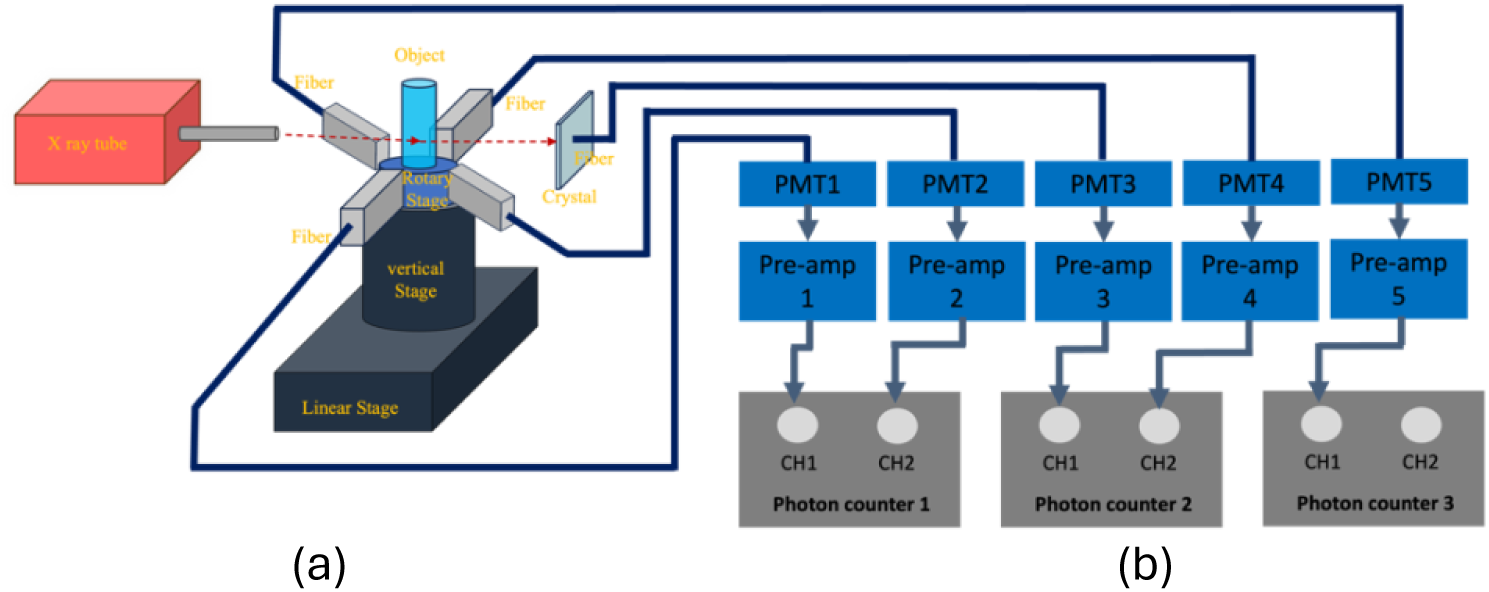
Schematic of XLCT imaging system with four detectors.

**Figure 2.**
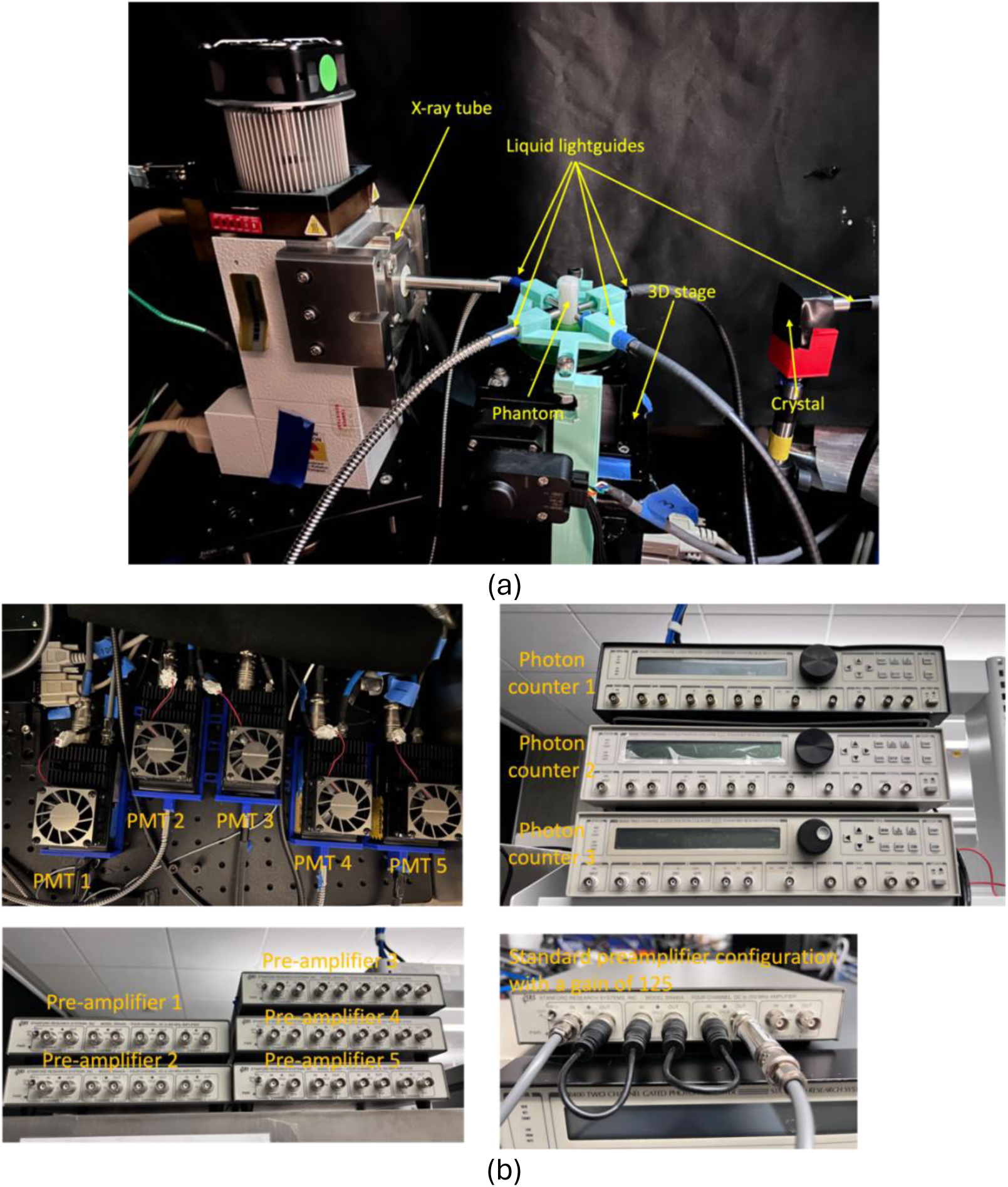
Photographs of the XLCT imaging system. (a) The x-ray tube with the x-ray optics lens, four fibers, and the single pixel x-ray detector; (b) 5 PMTs (left top); 3 photon counters (right top); 5 preamplifiers (left bottom); a typical connection of the preamplifier (right bottom).

To measure the x-ray luminescence signals for the XLCT imaging, we placed four liquid lightguides (fibers) around the object as shown in Fig 1. The angle between two close fibers is 90 degrees. The optical photons emitted from the object are gathered by fibers, each of which transmits the collected optical photons a PMT (H7422-50, Hamamatsu). As described above, the signals from the PMTs are then amplified using broadband amplifiers with a gain of 125. Subsequently, the signals are filtered and collected by the gated photon counters.

In the phantom experimental studies, we reconstructed the XLCT images with three different detector configurations: 1, 2, and 4 detectors. For the case of a single detector, measurements from one of the four detectors were selected. For the case of two detectors, measurements from two opposite (180 degrees apart) detectors were selected.

### 2.2 Weighted FBP algorithm

Traditional Filtered Back Projection (FBP) algorithm is widely used in imaging reconstruction of x-ray computed tomography (CT). FBP reconstructs CT images from measurements of attenuated x-ray intensity with x-ray detectors. In XLCT, we use x-ray beam to excite particles embedded inside tissue deeply and to measure the emitted optical photons from the particles on the object surface. We have to consider the x-ray beam attenuation and the optical photons’ scattering and absorption in the XLCT reconstruction.

In XLCT, the scanning pencil beam has the position information, and the detected optical photon number or intensity is proportional to the particle concentration and the x-ray beam density. Thus, a traditional FBP can reconstruct XLCT images with very good spatial resolution. The reconstructed XLCT images are only semi-quantitative because the scattering and the absorption of the optical photons are not considered, as shown by the following numerical simulations and phantom experiments. For most cases, semi-quantitative results are good enough, we can improve the results by including a sensitivity matrix of optical photons.

The optical photon propagation inside tissue can be calculated by the diffusion equation, from which we are able to obtain the sensitivity matrix ∑*P_d_* where d is detector number [19]. For each detector, we obtain one sensitivity matrix *P_d_* and then we add all the sensitivity matrixes as one matrix. The sensitivity matrix is usually based on a two-dimensional (2D) finite element matrix. The FBP reconstruction is performed on a 2D grid. We interpolate a new sensitivity matrix by interpolating the grids on the 2D finite element mesh.

The filtered back projection (FBP) algorithm involves filtering the raw projection data and then back projecting the filtered data to reconstruct the image. The equation can be expressed as:

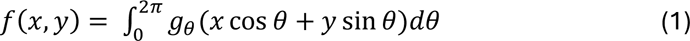

where *g_θ_* is the filter in the projection and *g_θ_* = ℎ(*r*) ∗ *p_θ_*(*r*). *p_θ_*(*r*) is the measurement of each projection *θ* and ℎ(*r*) is the filter.

The wFBP can be written as in Eq. (2). The weighting factor is the total sensitivity matrix as described above.

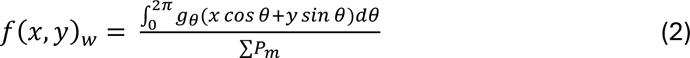

In this study, MATLAB was utilized for imaging reconstructions, employing both the FBP reconstruction and the wFBP reconstruction for all cases of XLCT imaging. The Hamming filter was employed in this study.

It is worth noting that we reconstruct the pencil beam CT images with the traditional FBP algorithm as described in Eq. (1).

### 2.3 Scan scheme of the XLCT imaging

During the experiment, we employed our laboratory-developed superfast scan scheme [Fang, IEEE access]. Briefly, Fig. 3 depicts the flowchart of the scan scheme. The photon counter consistently acquires pulse signals within a brief data collection window, typically set at 10 ms, and records position information at the window’s commencement for each measurement. The total measured pulse number is calculated by the specified scanning distance and speed of the linear stage. We scan one slice of the object by the specified angular projections and linear scan distance for each angular projection. Then, we scan multiple slices by adjusting the scanning height for 3D XLCT imaging. The space between each slice is adjustable by the step size of the vertical stage.

**Figure 3.**
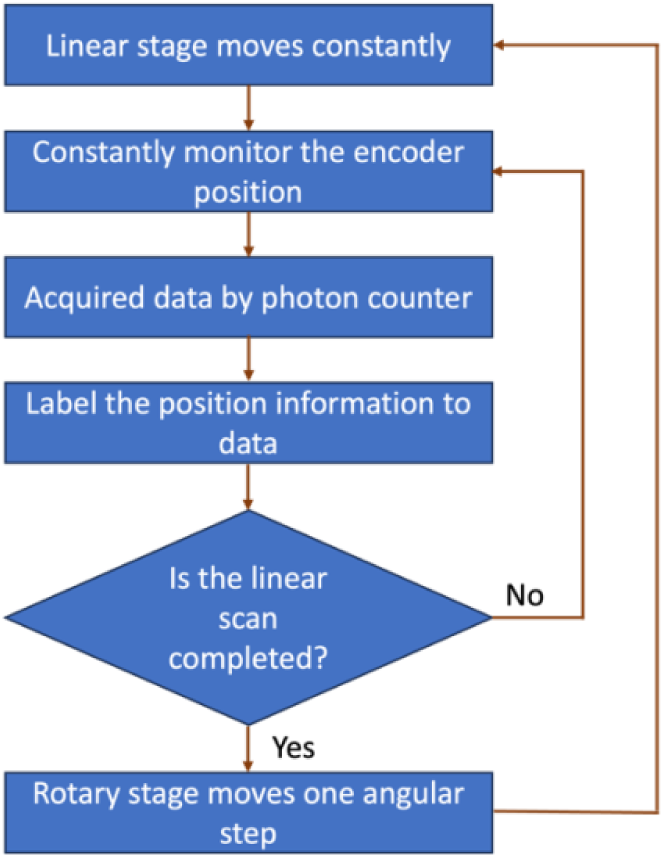
Flowchart of XLCT imaging system with a superfast scan scheme

### 2.4 Phantom experimental setup

Conducting phantom experiments involved utilizing a solid cylindrical agar phantom embedded with targets, and the schematic of the three different types of phantoms are shown in Fig. 4. The phantom’s background consisted of 1% intralipid, 2% agar, and water. We tested phantoms with two different sizes: 20 mm in height and 12 mm in diameter; 20 mm height and 20 mm in diameter. The embedded targets are glass capillary tubes filled with the solution of Gd_2_O_2_S: Eu^3+^ (GOS:Eu) (UKL63/UF-R1, Phosphor Technology. Ltd.) at concentrations of 10 mg/mL, 5 mg/mL, and 2.5 mg/mL, respectively.

**Figure 4.**
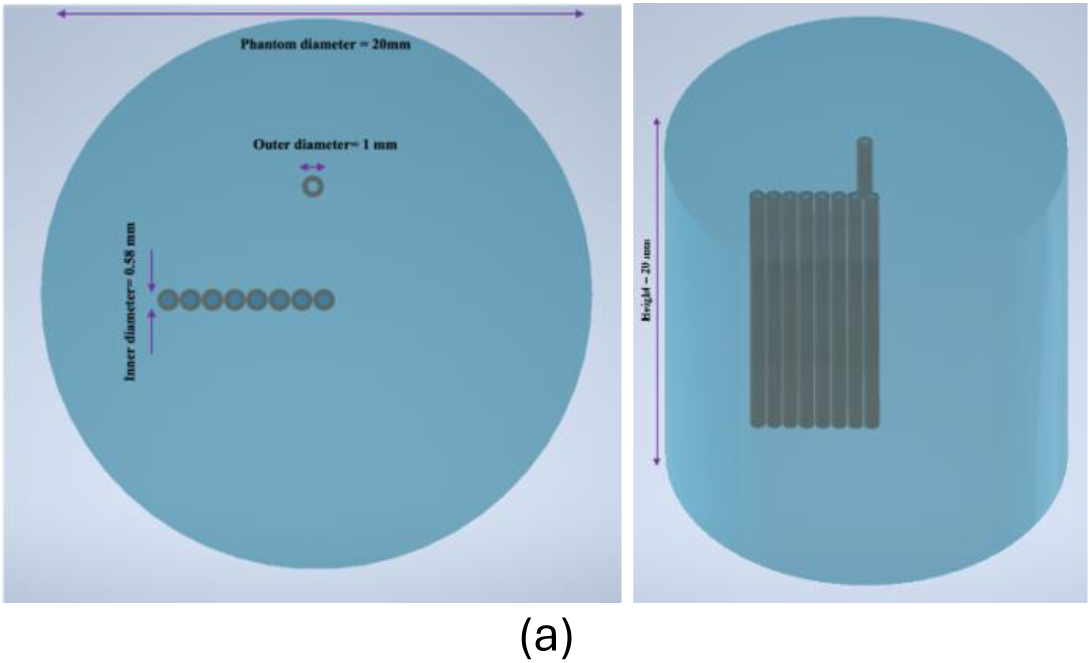
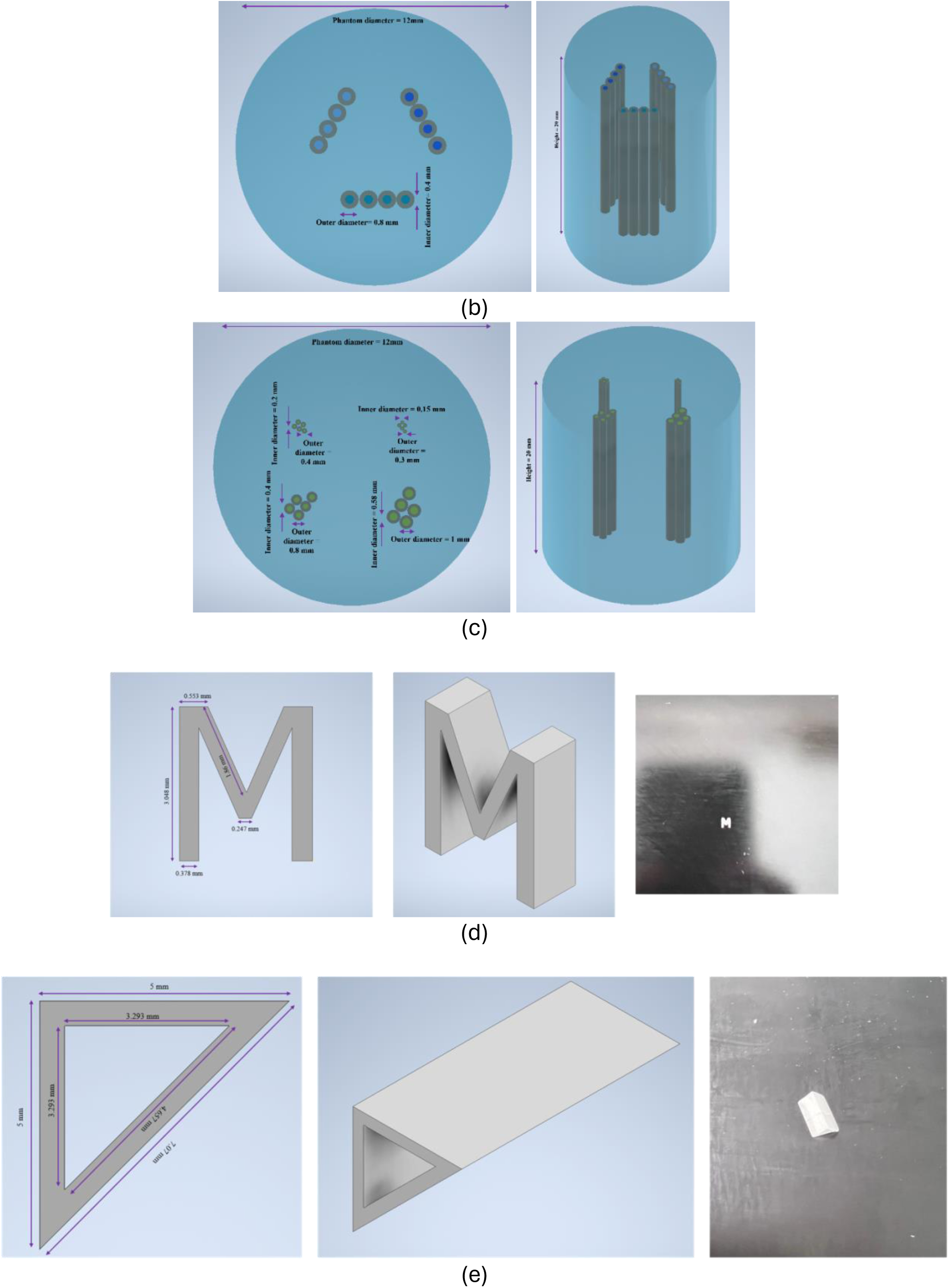
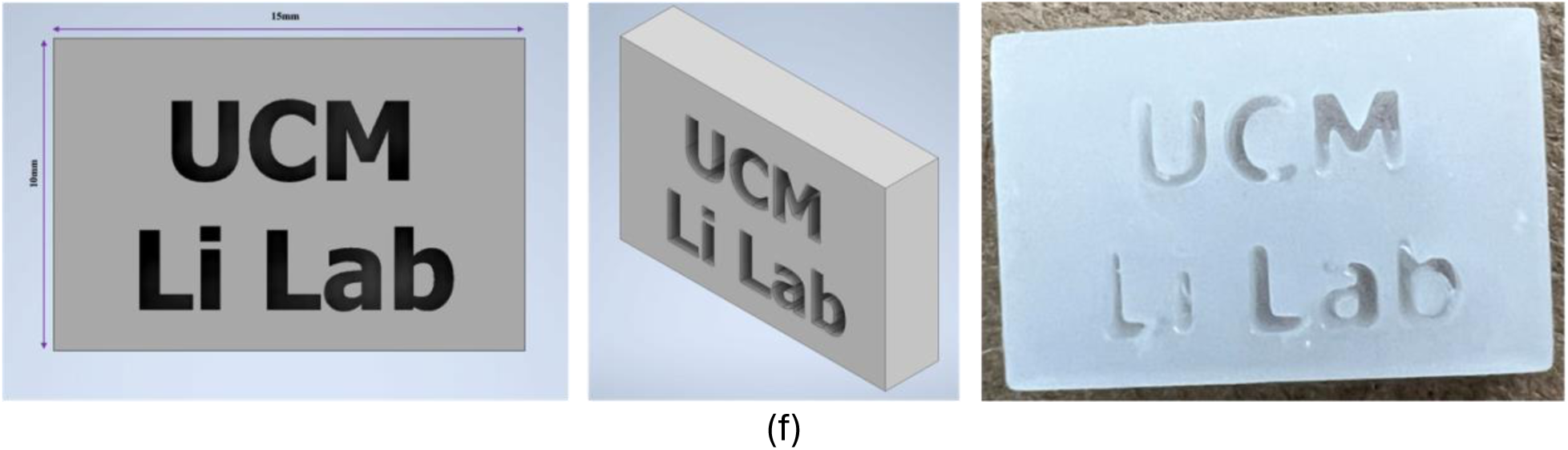
(a) Phantom with 8 targets aligned in a line (negative contrast phantom). (b) Phantom with 12 targets of three different concentrations. (c) Phantom with 20 targets of different sizes. (d) M target: (Left) front view with labeled dimension of the print target; (Middle) side view of the designed target; (Right) photo of the printed M target. (e) Triangular hollow bar target: (Left) the front view with labeled dimension of the print target; (Middle) the side view of the designed target; (Right) photo of the physical printed bar target. (f) The cuboid target with “UCM Li Lab” slot: (Left) front view with labeled dimension of the print target; (Middle) side view of the designed target; (Right) photo of the physical printed Cuboid target.

Fig. 4(a) shows the phantom for control contrast experiment with eight targets (0.58 mm of inner diameter and 1 mm of outer diameter) aligned of 10 mg/mL in a row and a single target (0.58 mm of inner diameter and 1.0 mm of outer diameter) with no particles above the row. The background of phantom has a 20 mm diameter. The aim of this experiment is to indicate that there is no scintillation from the glass capillary tubes.

Fig. 4(b) displays a phantom containing 12 targets with three different concentrations: 10 mg/mL, 5 mg/mL, and 2.5 mg/mL. Each group of targets with the same concentration has 4 targets. The capillary tube targets have an inner diameter of 0.4 mm and an outer diameter of 0.8 mm. The background of phantom has 12 mm diameter. This phantom is intended for the quantitative study of XLCT imaging.

Fig. 4(c) depicts four targets of different sizes, each filled with GOS solution at a concentration of 10 mg/mL. The inner diameters of the targets are 0.15 mm, 0.2 mm, 0.4 mm, and 0.58 mm, respectively. The outer diameters of the capillary tubes are 0.3 mm, 0.4 mm, 0.8 mm, and 1 mm. The background of the phantom has a diameter of 20 mm. This phantom is designed for spatial resolution study for the proposed system.

To explore the feasibility of the multichannel system, a unique 3D-printed target in the shape of a letter “M”, triangle hollow bar, and cuboid with “UCM Li Lab” slot were created. These distinctive targets were fabricated using clear resin blended with 10 mg/mL of GOS:Eu powder and produced through a 3D printer (ANYCUBIC Photon). The dimension of the “M” target is shown in Fig. 4 (d). The thickness of the printed “M” target is 1 mm thick. Fig. 4 (e) presents the dimensions of the triangle hollow bar along with a photograph of the printed bar. This target has an edge length of 5 mm and a thickness of 0.5 mm, with the hollow portion having an inner edge length of 3.293 mm. Fig. 4(f) depicts the dimensions of the cuboid target. Positioned at the center of the cuboid is a slot target in the shape of the “UCM Li Lab” logo. The cuboid measures 15 mm in length, 10 mm in width, and 3mm in height. During the experiments, the printed “M” target and the triangular hollow bar target were placed inside a 12 mm diameter agar phantom, and the cuboid target was placed inside a 20 mm diameter agar phantom.

### 2.5 Numerical simulation setup

To validate the spatial resolution and quantitative accuracy with the proposed wFBP algorithm, we have performed a numerical simulation using the traditional FBP and the wFBP reconstruction algorithms for measurement with 1, 2, 4, 8, and 16 detectors, respectively. For the numerical simulation, the phantom geometry is cylinder with a diameter of 12 mm and a height of 20 mm. The pixel size is 0.05 mm. We have 8 targets from the edge to the center of the phantom. The particle concentration of the 8 targets is set at 1 and the background has no particles. The diameter of the 8 targets is 0.4 mm and the center-to-center distance between two neighbor targets is 0.8 mm. The detectors are evenly distributed with an angular step of 360 degrees divided by the number of fibers. The angles between each fiber in cases with 1, 2, 4, 8, and 16 detectors are 360, 180, 90, 45, and 22.5 degrees, respectively. The numerical measurements were obtained following the method described in reference [20]. All the simulations were performed in MATLAB.

### 2.6 Evaluation Criteria

To evaluate the quantitative accuracy of the reconstructed XLCT images with FBP and wFBP methods, the ratio of reconstructed signals in targets were calculated. For this analysis, we selected the same region of interest (ROI) of each concentration within the same image slice and calculated the mean value of ROI. The dice coefficients were also calculated to evaluate the quality of the reconstructed XLCT images [20].

## 3. Results

### 3.1 Results of numerical simulation

Fig.5 plots the reconstructed XLCT images with 1, 2, 4, 8, and 16 detectors from left column to right column, respectively. The figures in the top row were reconstructed with FBP. The figures in the middle row were reconstructed with wFBP. The ground truth image is plotted in the bottom row of Fig. 5. Fig.6 plots show the line profiles across the eight targets for each reconstructed XLCT image. It is clear to see that the eight targets have been reconstructed successfully for different detector numbers. It is not surprising to see that the image quality is better with more detectors. Compared to the traditional FBP method, the wFBP reconstructed XLCT images with much better quality (less artifacts) and better quantitative accuracy as indicated by table 1 in which the reconstructed signal in each target was calculated and listed. From Figs. 5 and 6, we also see that the quantitative accuracy of deep targets (near phantom center) has been improved substantially with wFBP method, which is reasonable because deeper target has more scattering and absorption before reaching the phantom surface to be detected.

**Figure 5.**
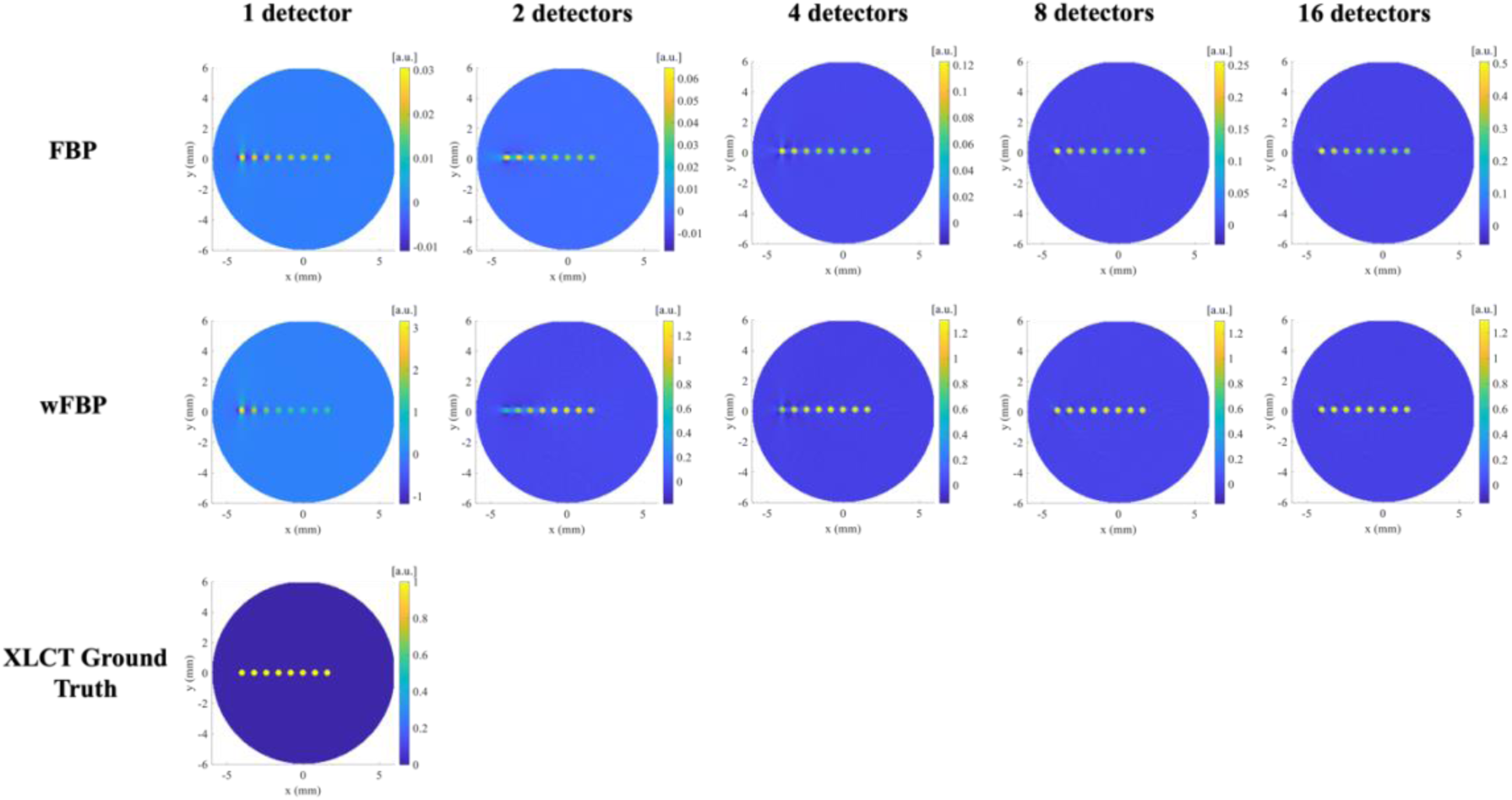
For the numerical simulation, the XLCT images were reconstructed with FBP (top row) and wFBP (middle row) and for different detector numbers. The bottom figure indicates the ground truth image.

**Figure 6.**
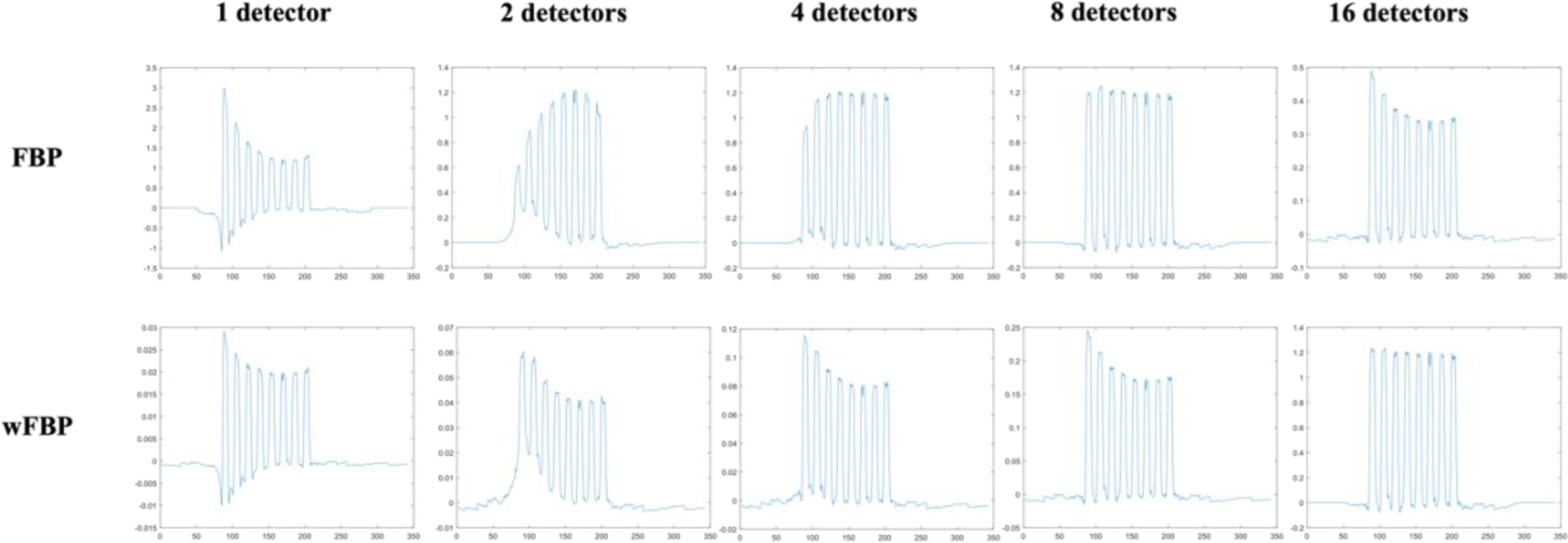
The line profile of phantom across eight targets for the reconstructed XLCT images in Fig. 5.

**Table 1.** Quantitative ratio of 8 targets for the numerical simulation.

| Detector numbers | wFBP quantitative ratio | FBP quantitative ratio |
| --- | --- | --- |
| 1 detector | 5.70 : 1.67 : 1 | 3.15 : 1.69 : 1 |
| 2 detectors | 3.21 : 1.64 : 1 | 3.15 : 1.69 : 1 |
| 4 detectors | 3.82 : 1.96 : 1 | 3.48 : 1.82 : 1 |
| 8 detectors | 3.79 : 1.94 : 1 | 3.48 : 1.82 : 1 |
| 816detectors | 3.79 : 1.95 : 1 | 3.48 : 1.82 : 1 |

### 3.2 Results of phantom experiments

#### 3.2.1 Control contrast phantom study

To verify that there is no scintillation signal from the glass capillary tube, we have performed a phantom experiment with a control contrast target (no GOS particle) and 8 GOS targets. The phantom geometry is plotted in Fig. 4(a). The XLCT images of this phantom were reconstructed with both FBP (top row in Fig. 7) and wFBP (bottom row in Fig. 7). We have also compared the reconstructed results from measurements with 1 detector (left column in Fig. 7), 2 detectors (middle column in Fig. 7) and 4 detectors (right column in Fig. 7). From Fig. 7, we see that all 8 GOS targets have been reconstructed successfully with both FBP and wFBP for different detector cases. We also see that the control target has no signals, which means that there is no scintillation from glass capillary tube indeed. We have also reconstructed the pencil beam CT images with FBP as plots in Fig. 8 (left).

**Figure 7.**
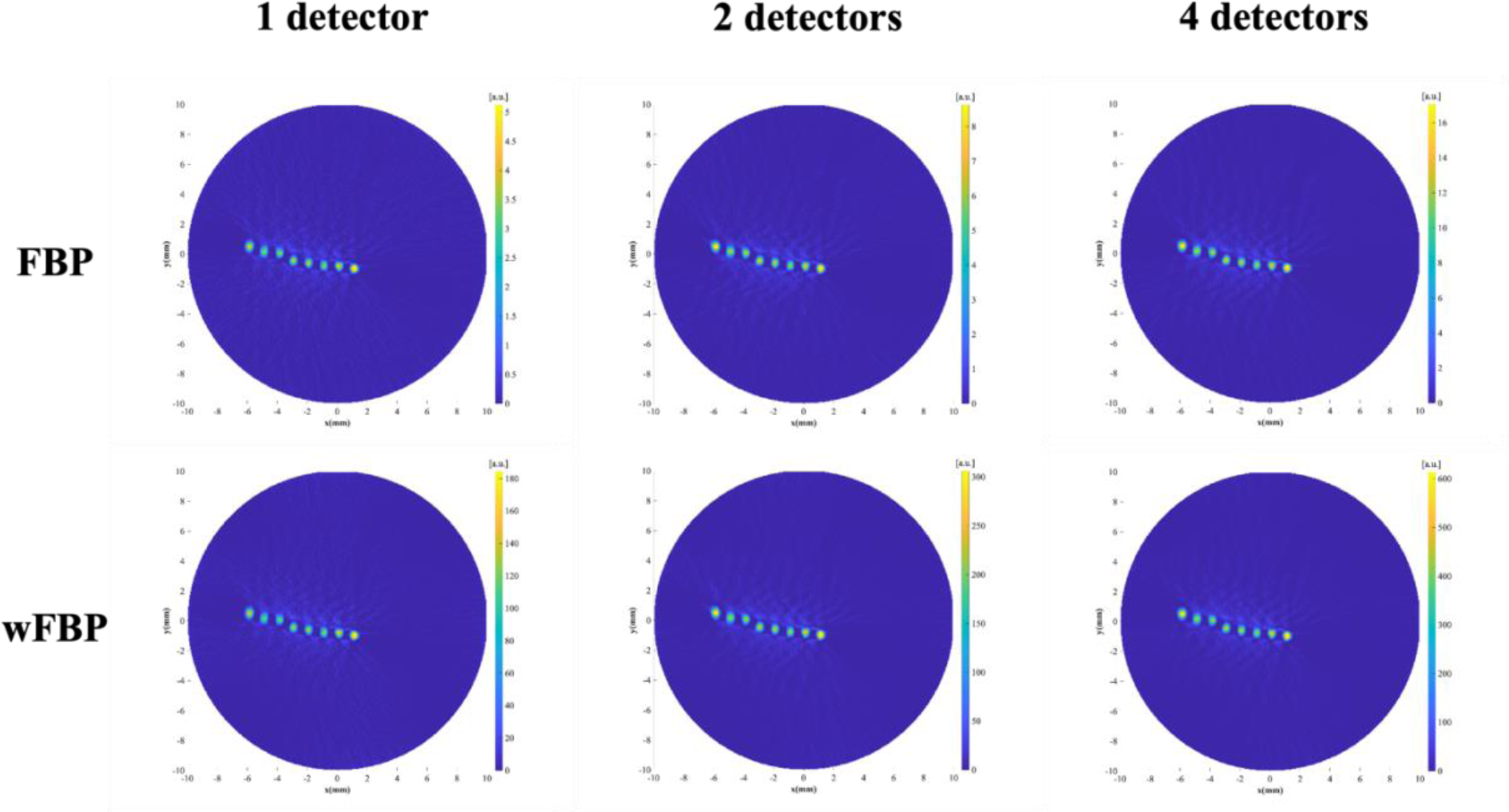
Reconstructed XLCT images of phantom with 8 GOS targets and 1 control target with FBP (top row) and wFBP (bottom row) for 1, 2 and 4 detectors, respectively.

**Figure 8.**
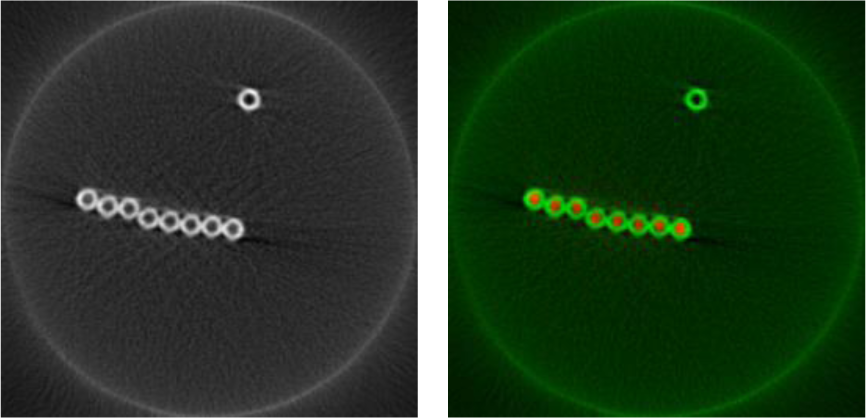
(left) Reconstructed pencil beam CT images. (right) The overlaid images of pencil beam CT (green) and the XLCT image reconstructed with wFBP and 4 detectors(color).

The reconstructed XLCT images for the case of 4 detectors and with wFBP is overlaid with the pencil beam CT images and is plotted in Fig. 8 (right). The reconstructed XLCT image is overlaid very well with the pencil CT image, which indicates that the XLCT image was reconstructed successfully in terms of target locations and sizes. Furthermore, it is observed that, after applying the wFBP method, the artifacts around the luminescent targets were reduced compared to the conventional FBP method. Additionally, in the middle of the 8 targets, some of the reconstructed targets have better contrasts using the wFBP method. To analyze the reconstructed XLCT images quantitatively, we have calculated the DICE coefficients of eight targets as shown in Table 2. From this table, we see that the DICE coefficients are improved slightly when the wFBP is applied for all detector cases.

**Table 2.**
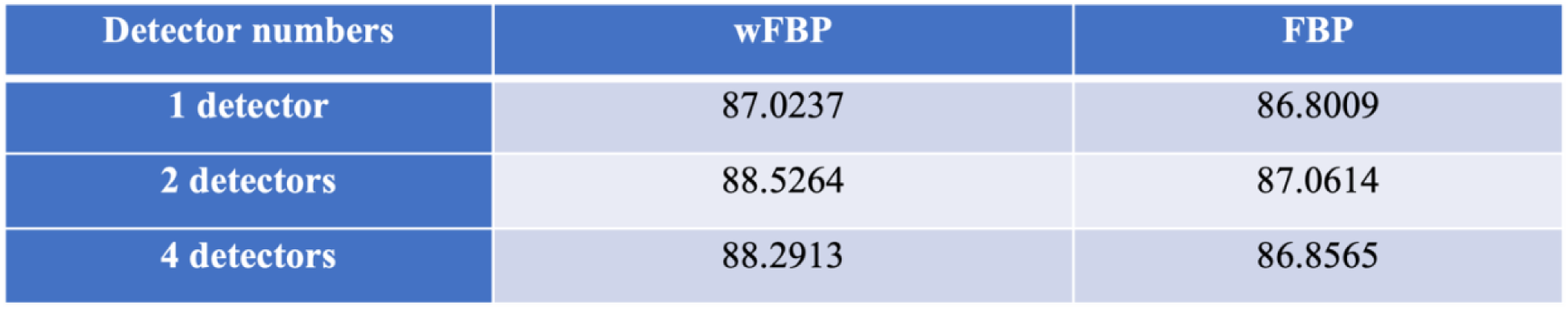
DICE coefficients of phantom with 8 GOS targets and 1 control target.

#### 3.2.2 Phantom experiment with different concentrations

Fig. 9 shows the reconstructed XLCT images of targets at different concentrations with the FBP and wFBP algorithms and using 1, 2, and 4 detectors, respectively. The left image in Fig. 10 plots the pencil beam CT images reconstructed with the FBP from the measurements of the single pixel crystal-based x-ray detector. The pencil beam CT can only detect the capillary tubes and the agar background phantom and is unable to visualize the luminescent targets due to their low concentration. We have overlaid the XLCT images (the bottom right figure in Fig. 9) with the pencil beam CT images as shown in Fig. 10 (right), from which we see that the pencil beam CT targets (green circles) align very well with the XLCT targets (red dots).

**Figure 9.**
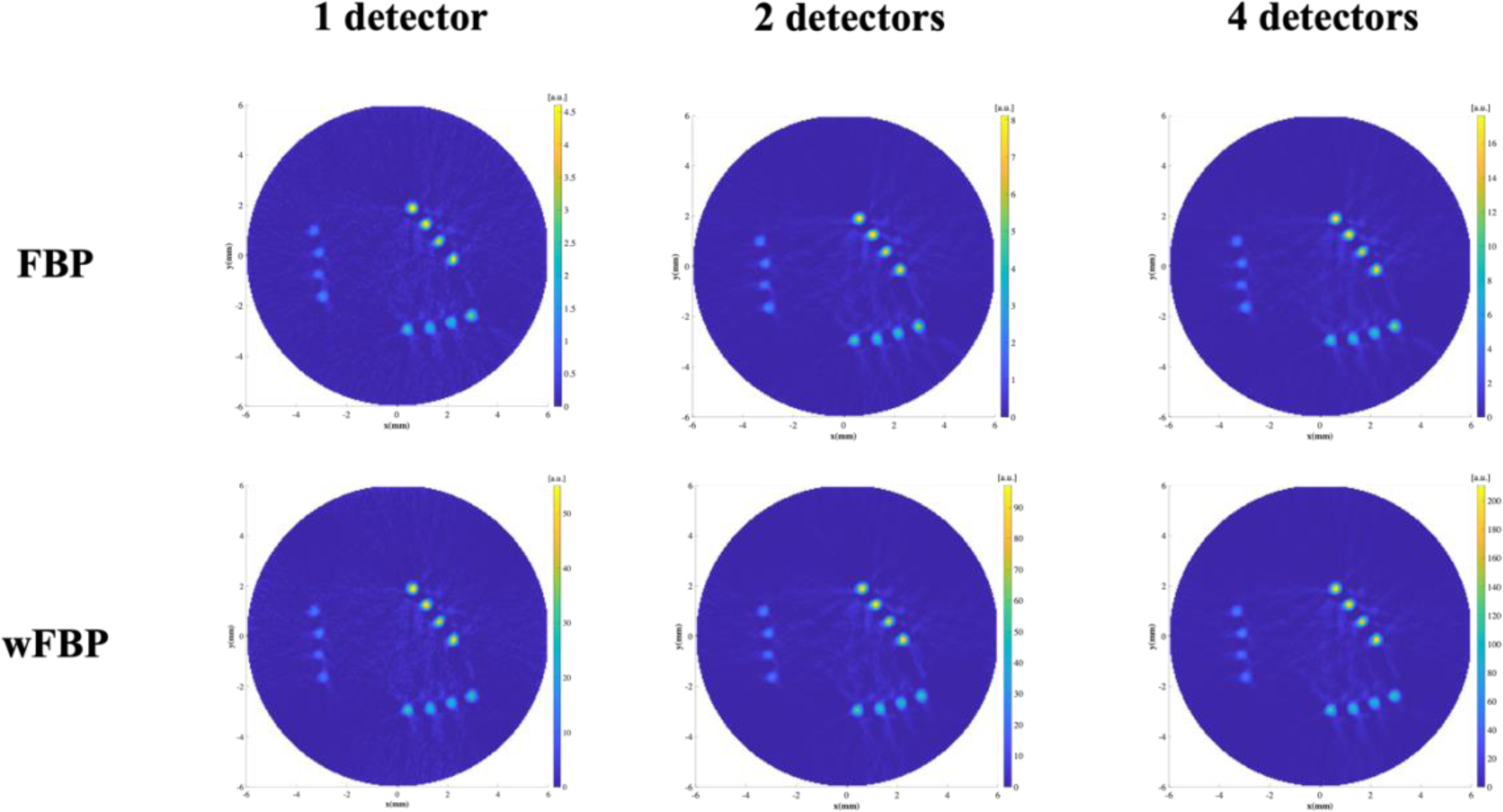
Reconstructed XLCT images for the phantom with three different concentrations in 12 targets for FBP (top row) and wFBP (bottom row), and for cases of 1 detector (left column), 2 detectors (middle column), and 4 detectors (right column)

**Figure 10.**
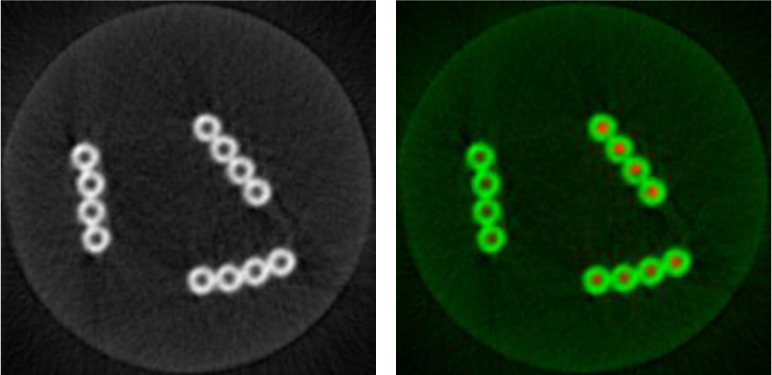
(left) Reconstructed pencil beam CT images of the phantom with different concentrations. (right) The overlaid images of pencil beam CT (green) and the XLCT image reconstructed with wFBP and 4 detectors(color).

Both XLCT images and parallel pencil beam CT images were successfully reconstructed, as shown in Fig.9 and Fig. 10. The 12 capillary tubes in the pencil beam CT image were accurately reconstructed and easily observed. The circular light gray shape corresponds to the agar background of the phantom. The reconstructed image of capillary tube targets from pencil beam CT serves as the ground truth, offering readily available position information for the targets. Difference intensity of each set of concentration targets are obverse to show in the reconstructed XLCT images from 1, 2, and 4 detectors configurations. The yellow, light blue, and dark blue colors of the group of targets in the images indicate the different measurement intensity of the targets. As depicted in Fig. 9, the reconstructed images of XLCT targets with a 4-detector configuration exhibit have less artifacts compared to the images reconstructed with either 2 detectors or 1 detector configurations. This suggests that the use of 4-detector configuration achieved the best image quality. We can also see that the reconstructed XLCT images with the wFBP have less artifacts than the XLCT images with the traditional FBP.

The data for both pencil beam CT images and luminescence CT images were acquired simultaneously. The angular projection is 360 with an angular step size of 1 degree.

Table 3 shows the quantitative ratios of reconstructed images for the phantom with different concentrations using setups with 1, 2, and 4 detectors. We selected the same area around each set of 4 targets with identical concentrations. The mean value of the selected area was then calculated, followed by determining the ratio of each concentration using both the wFBP and the FBP algorithms. It is evident that the ratios obtained through the wFBP reconstruction method are closer to the true ratios of target concentrations (10 mg/mL:5 mg/mL:2.5 mg/mL) than the traditional FBP method when employing 1, 2, and 4 detectors. This indicates that the proposed wFBP can improve the target signals quantitatively.

**Table 3.** Quantitative ratios of phantom with three different concentrations.

| Detector numbers | wFBP quantitative ratio | FBP quantitative ratio |
| --- | --- | --- |
| 1 detector | 3.86 : 1.75 : 1 | 3.58 : 2.1 : 1 |
| 2 detectors | 3.74 : 1.8 : 1 | 3.53 : 2.1 : 1 |
| 4 detectors | 3.8 : 1.79 : 1 | 3.42 : 2 : 1 |

Table 4 shows the DICE similarity coefficients of each case to quantitively analysis the accuracy of the image reconstruction. For the FBP algorithm, we calculated the DICE are calculated to be 79.9%, 80.2%, and 81.4% with 1 detector, 2 detectors, and 4 detectors, respectively. For the corrected FBP algorithm, we calculated the DICE are calculated to be 79.3%, 80.1%, and 80.8% with 1 detector, 2 detectors, and 4 detectors, respectively. We can see that both algorithms have a high similarity between the reconstructed targets and the ground truth. And the setups with more detectors achieved slightly better DICE coefficients, which is as expected because of better efficiency of collecting photons from different directions.

**Table 4.** DICE coefficients of phantom with different concentration.

| Detector numbers | wFBP | FBP |
| --- | --- | --- |
| 1 detector | 79.3057 | 79.9191 |
| 2 detectors | 80.0808 | 80.2428 |
| 4 detectors | 80.7588 | 81.3652 |

#### 3.2.3 Phantom experiments with different target sizes

The purpose of scanning this phantom is to investigate the high spatial resolution imaging capabilities of the multichannel-based XLCT system. Fig. 11 displays the reconstructed images of a phantom with different target sizes. Both pencil beam CT and XLCT images were reconstructed for this phantom. The XLCT targets have diameters of 0.15 mm, 0.2 mm, 0.4 mm, and 0.58 mm, respectively. Each target group has 5 targets. The XLCT images were reconstructed with both FBP and wFBP for the detector cases (1, 2, and 4 detectors).

**Figure 11.**
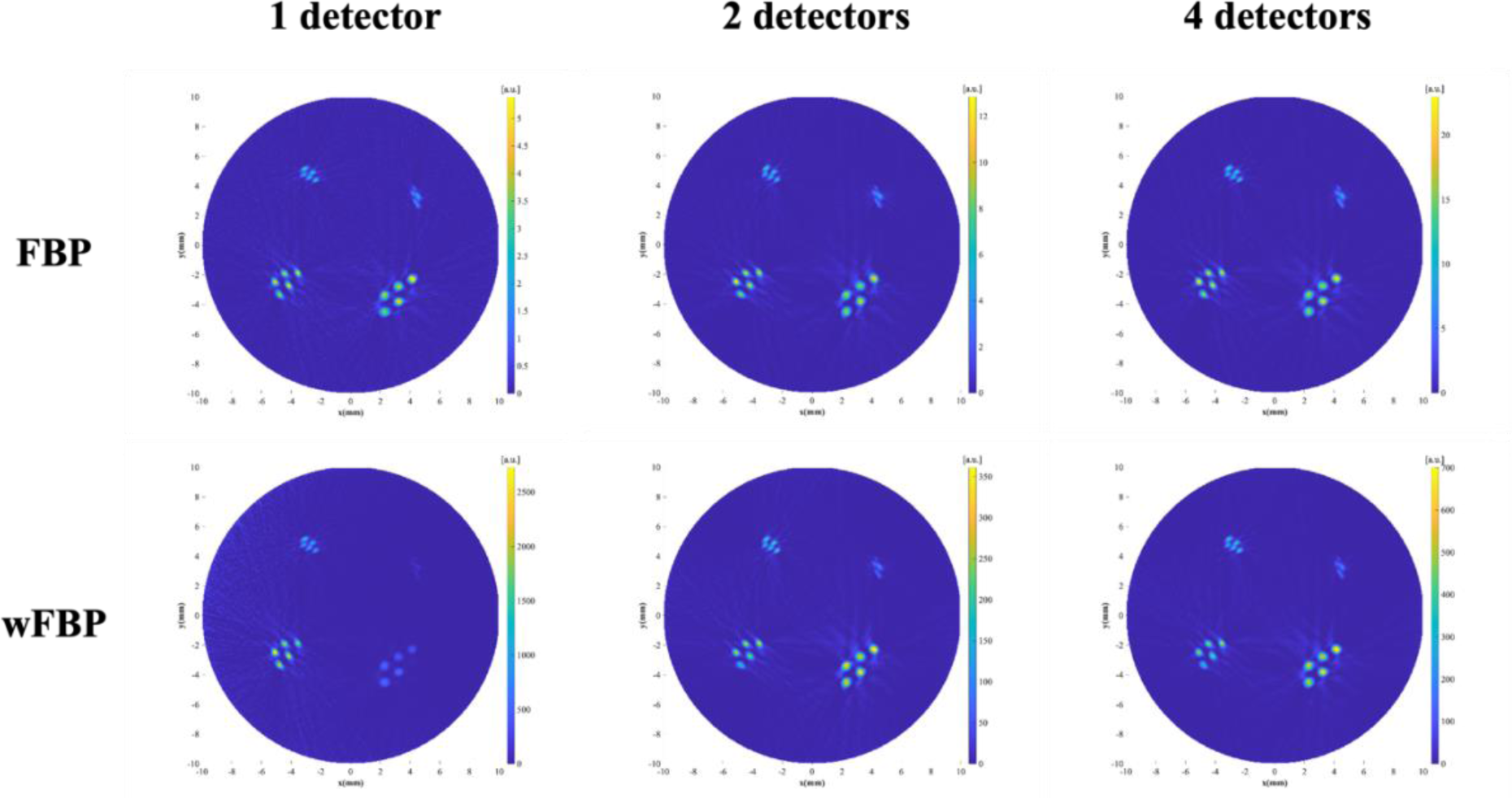
Reconstructed XLCT images for the phantom with four different sizes for FBP (top row) and wFBP (bottom row), and for cases of 1 detector (left column), 2 detectors (middle column), and 4 detectors (right column)

As depicted in Fig. 12, the position of the capillary tube targets was successfully reconstructed by the pencil beam CT, providing a clear view of the four groups of targets with different sizes. The capillary tube targets with diameters of 1 mm, 0.8 mm and 0.4 mm have been successfully and easily differentiated in the reconstructed XLCT images. The targets in the group with an outer diameter of 0.3 mm and inner diameter of 0.15 mm were also differentiated but marginally. And in the overlaid image, we can see the CT targets are overlaid very well with the XLCT targets like the previous phantom experiment. To analyze the reconstructed XLCT images quantitatively, we calculated the DICE coefficients for 20 targets, as shown in Table 5. The table indicates that the DICE coefficients improved slightly when the wFBP method was applied across all detector cases.

**Figure 12.**
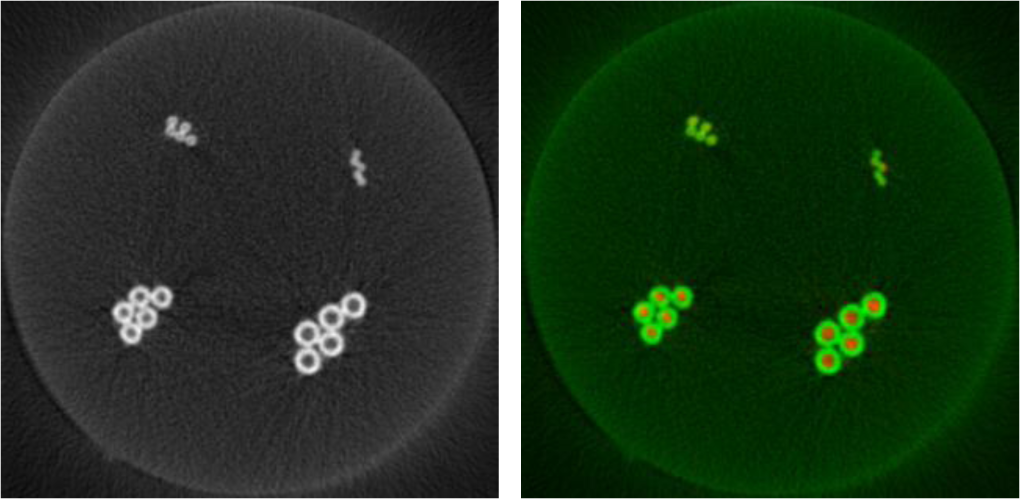
(left) Reconstructed pencil beam CT images of the phantom with 4 different sizes. (right) The overlaid images of pencil beam CT (green) and the XLCT image reconstructed with wFBP and 4 detectors.

**Table 5.**
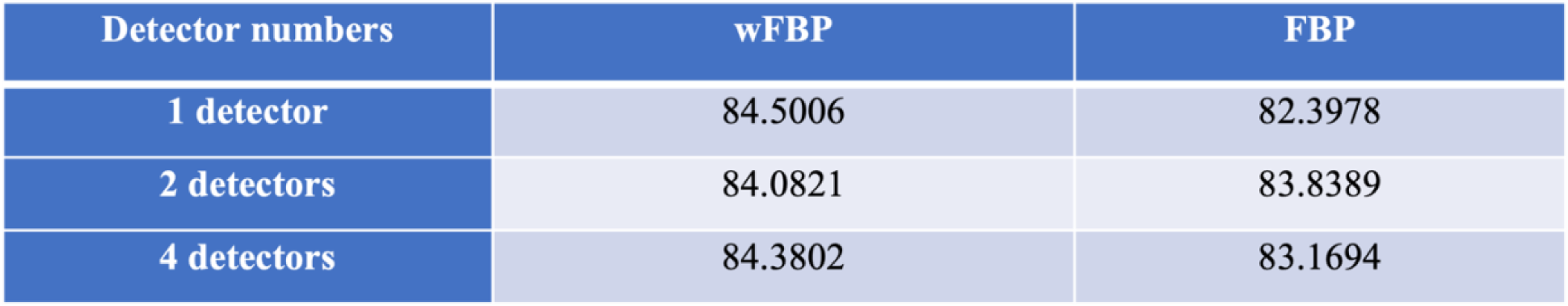
DICE coefficients of phantom with different size.

| Detector numbers | wFBP | FBP |
| --- | --- | --- |
| 1 detector | 84.5006 | 82.3978 |
| 2 detectors | 84.0821 | 83.8389 |
| 4 detectors | 84.3802 | 83.1694 |

#### 3.2.4 Phantom experiments of 3D XLCT imaging

In the experiment of M target, we scanned 90 angular projections for each transverse scan with an angular step size of 4 degrees. To create 3D images, we conducted a total of 11 transverse sections with a vertical step size of 0.2 mm. The thickness of M target is 1 mm. Because the target was not perfectly placed horizontally in the background phantom, a vertical scan of 2 mm covered the entire M target. The results of the reconstructed images are shown in Fig. 13 for this experiment. In Fig. 13 (a), the reconstructed 3D XLCT images with both the wFBP and FBP methods are plotted, which has shown that with an increase in the number of detectors, more photons were collected for imaging reconstruction and significant reduction of artifacts around the M target was achieved. The application of the wFBP further reduced artifacts around the target M, particularly for the case with only 1 detector.

**Figure 13.**
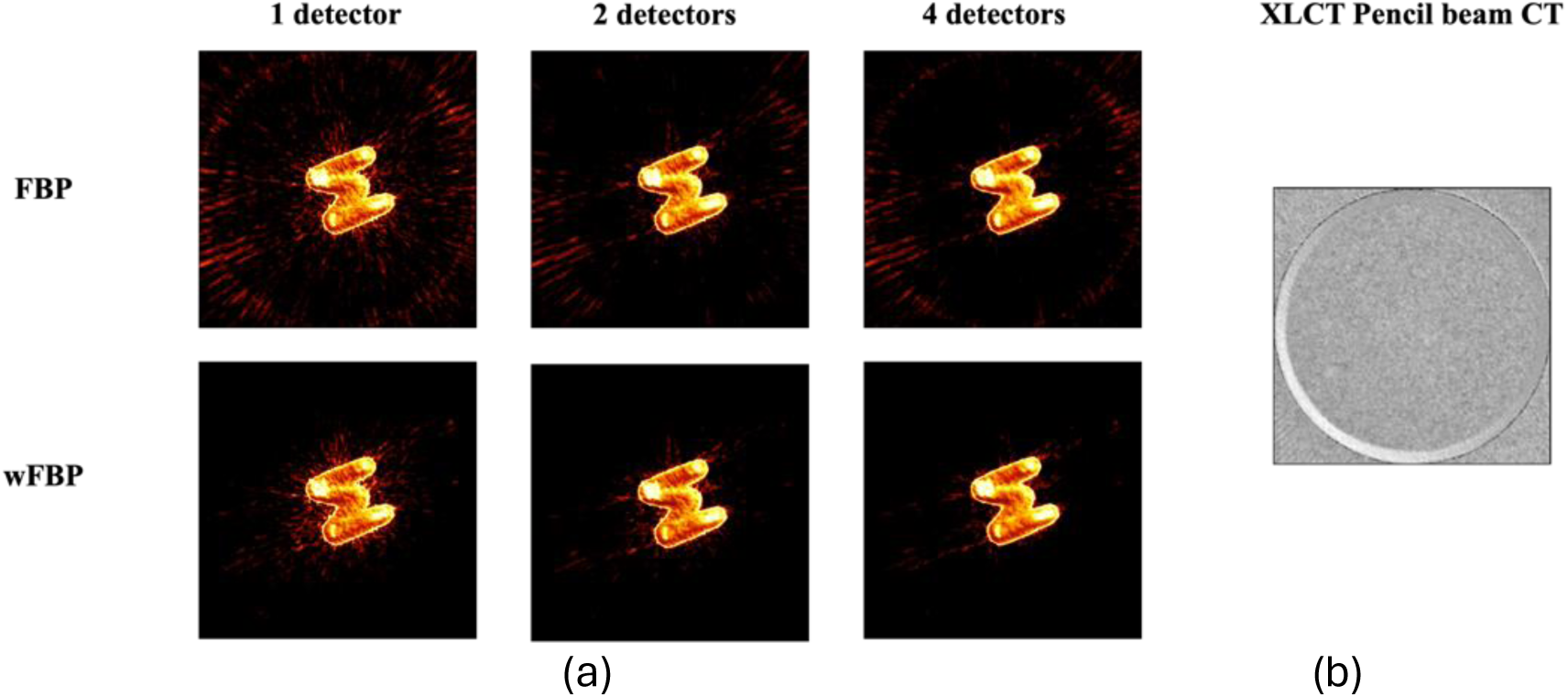
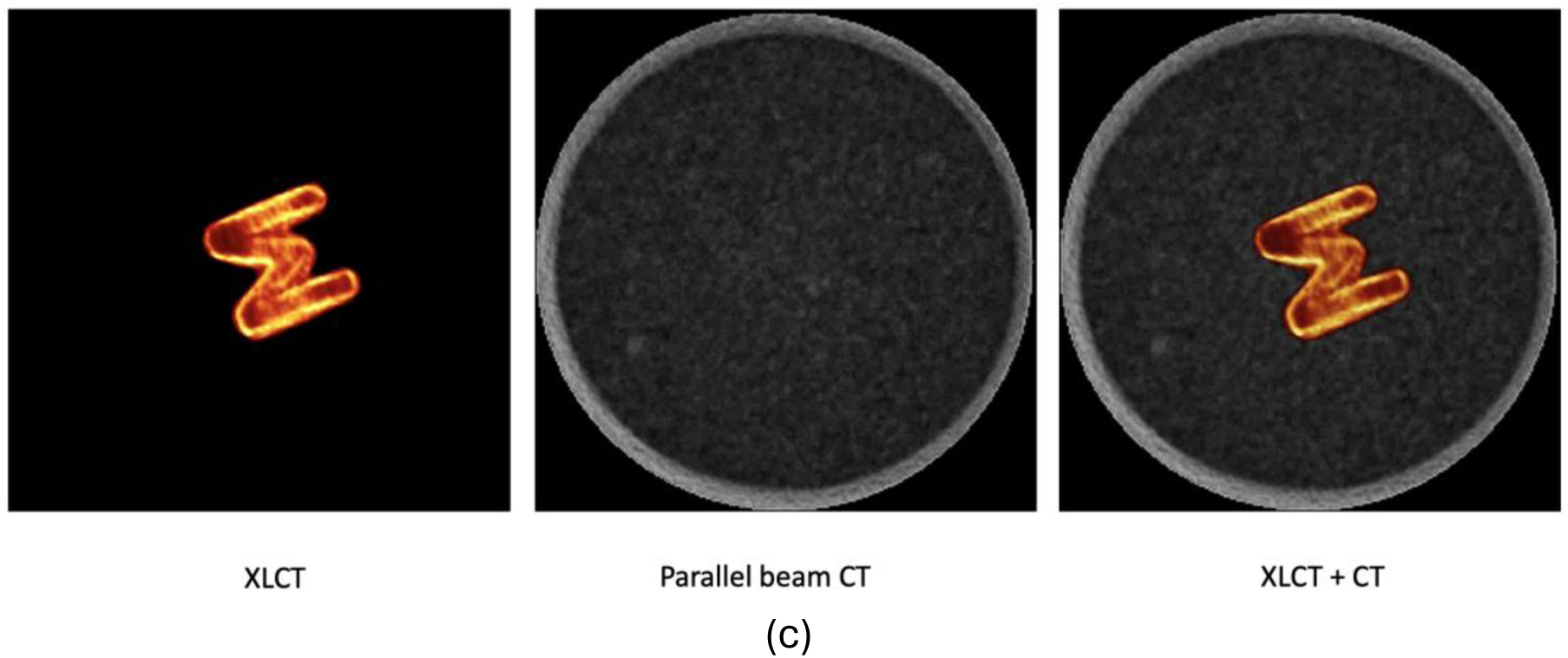
The experimental results of the “M” phantom. (a) Reconstructed 3D XLCT images with detector numbers of 1 (left), 2 (middle), and 4 (right), and with the reconstruction method of FBP (top) and wFBP (bottom). (b) The reconstructed pencil beam CT images. (c) The rendered 3D movies of the reconstructed XLCT images (left), pencil beam CT images (middle), and overlaid images (right). Movies are shown in the supplemental media.

The reconstructed pencil beam CT images are plotted on the right side of Fig. 13(b), in which we can only see the background of the phantom and the M target is not visible as expected. Fig. 13(c) depicts the rendered 3D image movie (see supplemental media) of the reconstructed XLCT (left), pencil beam CT (middle), and the overlaid images of XLCT and pencil beam CT (right) for the case with 4 detector and the wFBP method. The rendered 3D images were formed from the 11 transverse images.

Notably, the M target was manufactured using a 3D printer, and the material used is resin mixed with Gd₂O₂S:Eu³⁺ (GOS:Eu) powder. During the pencil beam CT imaging reconstruction, only the agar background is visible in the pencil beam CT images due to the smaller quantity of resin used.

We have also conducted two additional experiments using a triangular hollow bar target and a cuboid with a “UCM Li Lab” slot target. During the experiments, the triangular hollow bar target was placed inside a 12 mm diameter agar phantom, and the cuboid target was placed inside a 20 mm diameter agar phantom. The production process of these two phantoms was the same as that of the capillary tube phantoms. These two experiments evaluated the performance of the XLCT system in imaging hollow targets and hot background targets.

Fig. 14(a) plots the reconstructed 3D images of the triangular hollow bar phantom: XLCT images with the wFBP and 4 detectors (left), pencil beam CT images with FBP method (middle), and their overlaid images (right). From this figure, we see that the hollow bar target has been reconstructed successfully. In the experiment, we scanned 90 angular projections for each transverse scan with an angular step size of 4 degrees. To create 3D XLCT and pencil beam CT images, we conducted a total of 6 transverse sections with a vertical step size of 0.5 mm.

**Figure 14.**
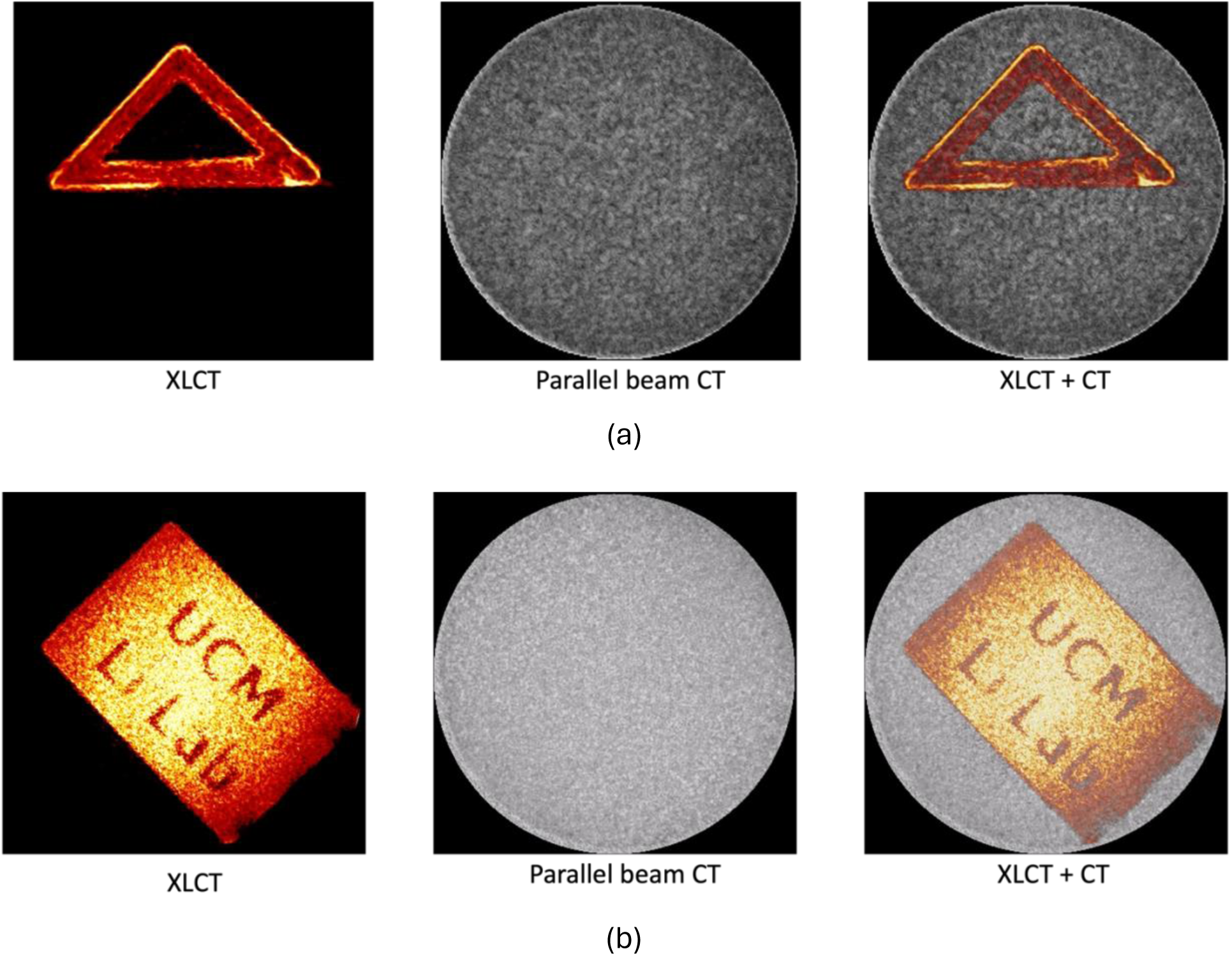
The reconstructed 3D images of XLCT and pencil beam CT for the hollow triangle target (a) and the cuboid target (b): Reconstructed 3D XLCT images with 4 detectors and wFBP (left); the reconstructed pencil beam CT images (middle), the overlaid images (right). The movies are shown in the supplemental media.

Fig. 14(b) shows the reconstructed 3D imaging of the cuboid target with “UCM Li Lab” slot. The left, middle, and right images show the reconstructed XLCT image with the wFBP and four detectors, parallel pencil beam CT with FBP, and their overlaid images, respectively. For each transvers section, we scanned 90 angular projections with the 4- degree angular step size. To create the 3D image, we scan 11 layers, and the vertical step size is 0.5 mm.

Notably, reconstructing the cuboid target with the “UCM Li Lab” target is very challenging due to the tiny and irregular shape of the target and to the hot background. Nevertheless, we successfully reconstructed it using our XLCT system with the wFBP reconstruction method. In the left image of Fig. 14(b), the edge of the cuboid appears slightly darker than other areas of the target because the target edge is closer to the edge of the phantom so that the attenuation correction is harder. Additionally, due to the 3D printer’s resolution limitations, the printed slot in the cuboid is not as accurate as we designed. The dot on the “i” was missing because of this printer low resolution issue. Despite these drawbacks, we successfully reconstructed this very complex target, demonstrating that our XLCT system is capable of imaging targets with very complex and irregular shapes and with a hot background.

## 3 Discussion and conclusion

In this work, we have, for the first time, introduced and experimentally demonstrated a multichannel-based X-ray Luminescence Computed Tomography (XLCT) system, the number of detectors up to 4. The weighted Filtered Back Projection (FBP) algorithm was, also for the first time, applied to the reconstruction of XLCT images at very high spatial resolution.

The proposed multichannel XLCT system and the weighted FBP reconstruction algorithm were studied with 1, 2, and 4 detectors. And all the results of the reconstructed images were compared with the traditional FBP method. To explore the performance of our system, we perform numerical simulation studies and physical experimental studies. Due to the limitation of our current equipment, the maximum number of detectors are chosen to be 4. In the numerical simulation, we simulated the cases up to 16 detectors. In the numerical investigations, an increase in the number of detectors resulted in better reconstructions of the XLCT targets, as expected. Fig. 5 provides the best explanation of this trend. In the case of a single detector, several artifacts around the targets are observed since only a portion of the scatter-excited photons was collected. As the number of detectors increased, the artifacts around the targets significantly reduced, resulting in clearer reconstructed targets. This improvement is attributed to the collection of more optical photons by the detectors. Table 1 displays the quantitative ratio of targets with varying concentrations obtained from both algorithms, confirming that the wFBP algorithm results in a more accurate ratio than the FBP.

Following the numerical simulation results, we conducted physical experimental studies to evaluate the performance of the multichannel XLCT system. Three types of phantoms (different targets sizes, different concentrations, and control contrast) were created to evaluate the XLCT imaging system’s spatial resolution, and quantitative accuracy, respectively. All the phantoms were scanned by 1, 2, and 4 detectors with 360 angular projections with an angular step of 1-degree. The spatial resolution study results are presented in Fig. 11. The XLCT imaging system can successfully reconstruct minimum targets, an edge-to-edge distance of 0.15 mm, which is close to the x-ray beam size of 0.15 mm. In future experiments, a super-fine beam x-ray tube will be employed to achieve even smaller minimum resolutions. The quantitative study results are shown in Fig. 10 and table 3. Table 3 quantifies the concentration ratios, highlighting the superior accuracy of the wFBP algorithm compared to the traditional FBP method, thereby emphasizing its potential for precise and reliable quantitative analyses in medical imaging and diagnostics. The control contrast imaging study validated that there is no scintillation from the capillary tube.

To evaluate the XLCT system’s capability for 3D imaging, we scanned three different targets: a M target, a triangle hollow bar, and a hot background cuboid with “UCM Li Lab” slot target. The targets were manufactured using resin mixed with Gd₂O₂S:Eu³⁺ (GOS:Eu) powder. The experiment of the M target employed 1, 2, and 4 detector configurations, utilizing both the wFBP and the traditional FBP algorithms. The results, displayed in Fig. 13 (a). Fig. 13(c), exhibited reduced artifacts around the 3D printed targets with an increase of detector number. All these targets have been reconstructed successfully, particularly for the case of the hot background cuboid, demonstrating the power of the XLCT imaging system and the proposed weighted FBP method.

In sum, both the numerical simulations and phantom experiments have demonstrated the feasibility and the efficacy of the multiple channel XLCT imaging system. In the future, further refinements and optimizations, especially in addressing challenges related to target size and positioning, could contribute to the continued advancement of XLCT as a valuable imaging modality in preclinical research.

## Supporting information

Supplemental Figure 13, and will be used for the link to the file on the preprint site.

Supplemental Figure 14(a), and will be used for the link to the file on the preprint site.

Supplemental Figure 14(b), and will be used for the link to the file on the preprint site.

## Notes

### Competing Interest Statement

The authors have declared no competing interest.

